# Changes in lactic acid concentrations in culture media and its implications in the inhibition of fungal development

**DOI:** 10.1101/2023.08.10.552543

**Authors:** J. Hinojosa Laso

## Abstract

In the following research, a novel antifungal culture method was discussed with the aim of developing bacterial culture media in which fungal growth is inhibited and bacterial growth is stimulated. This technique is based on the use of lactic acid as an antifungal agent, due to its capacity to neutralise the electrochemical gradient of fungal cell membranes, thus inhibiting their development. Based on this principle, the Minimum Inhibitory Concentration (MIC) of lactic acid in the culture media was investigated by changing its concentrations. After the preparation of the solutions, microbiological seeding was carried out in the different experimental groups. Data were collected using a quadrat sampling technique for subsequent statistical analysis using the T-test coefficient. As a result, the minimum inhibitory concentration of lactic acid was found to be 20%, where fungal growth was inhibited, and bacterial growth was stimulated.

**RESUMEN:** En la presente investigación, se discutió acerca de un novedoso método de cultivo antifúngico con la finalidad de desarrollar medios de cultivo bacterianos; en los que el crecimiento de hongos se viera inhibido y el bacteriano estimulado. Esta técnica se basa en el uso del ácido láctico como agente antifúngico, debido a la capacidad de este ácido para neutralizar el gradiente electroquímico de las membranas celulares de los hongos, inhibiendo así su desarrollo. A partir de esta premisa, se investigó la Concentración Inhibitoria Mínima (CIM) del ácido láctico en los medios de cultivo mediante un cambio en sus concentraciones. Posterior a la preparación de las soluciones, se procedió a la siembra microbiológica en los diferentes grupos experimentales. Los datos fueron recolectados mediante una técnica de muestreo por cuadrantes para su posterior análisis estadístico mediante el coeficiente de la Prueba T. Como resultado, se encontró que la concentración inhibitoria mínima del ácido láctico fue del 20%, donde el desarrollo fúngico se vio inhibido y el bacteriano estimulado.

**Published Electronically:**

- *Background and Aims* Lactic acid has been shown to have a negative effect on the chemical activity of fungal cell membranes and is therefore proposed to be used as an antifungal agent in growth media.
- *Methods* In the current study, an 88% lactic acid solution was used to produce agar solutions for culture media to assess the extent to which lactic acid can inhibit fungal growth. Subsequently, a quadrant sampling technique and a statistical study were used to measure the extent to which lactic acid can inhibit the growth of fungi.
- *Key Results* An inversely proportional correlation was found where lactic acid inhibited fungal growth. Specifically, the minimum inhibitory concentration of lactic acid was 20% and the results were supported by mathematical correlation and a statistical T-test, where 95% reliability was assumed.
- *Conclusions* These results are consistent with the limited research into the molecular pathways of lactic acid in fungal organisms, which establishes, as in this research, that lactic acid negatively affects fungi and their development in culture media.

## INTRODUCTION

Microbiology is the field of biology that oversees studying microorganisms, including bacteria and fungi, in a holistic way by also investigating their relationships with the biotic and abiotic environment (Universidad de los Andes, n.d.). Due to the nature of this discipline, its study and experimentation are usually carried out using culture media.

In its definition, culture media are substrates or solutions with which it is sought to stimulate the proliferation of microorganisms, such as bacteria or fungi, for their subsequent study. To achieve this, it is necessary that these solutions be enriched; therefore, they must contain nutrients, often sugars, agar, or nutrient broths, which serve as food for the microorganisms and as stimulators of their proliferation (Gómez, G., Batista, C., 2006).

Thanks to the various components that can be added to these media, the production of culture media for different purposes can be achieved. Firstly, there are selective media, which are used to stimulate or favour the growth of a certain species within a bacterial community. There are also inhibitory media, which contain certain substances that prevent the appearance, growth, and reproduction of a certain species or type of microorganism (Barona, 2021, slides 7-8).

These latter media are of great importance, since, generally, when an experiment is conducted, the development and reproduction of a single type of microorganism is pursued, and the contamination of this could have a negative effect on the growth of the desired micro-organisms. An example of this, and what will be investigated in this paper, is the development of a culture medium that can inhibit only fungal growth and not bacterial growth.

For this, it would be necessary to have or develop an agent or factor that, at a specific concentration, also known as minimum inhibitory concentration (MIC), inhibits fungal growth and development, but not bacterial growth (Dong, L., et al., 2012). These factors or agents are known as antifungal agents, which have a high specificity, acting only on the metabolic pathways or unique structures of fungi, being beneficial since they would only prevent the development of fungal organisms and not other types of microorganisms, such as bacteria (Susana, B., 2005); thus, allowing the development of antifungal culture media.

For the development of these culture media, lactic acid is proposed, which is obtained as a by-product of lactic fermentation, and its application in experimentation is novel since, as will be mentioned below, the molecular mechanisms of this agent have not been explored in depth (Vélez, C., León, A., 2014).

### Previous Research

As previously stated, the use of lactic acid as an antifungal agent is a novel application, so it was only possible to find a review of this technique carried out by the University of Antioquia, in Colombia. In this review, it was concluded that lactic acid, in synergy with acetic acid, forms an interaction with the fungal cell membrane that produces a neutralisation of the electrochemical proton gradient, thus affecting cell transport, and an inhibition in the uptake of amino acids by the cell structure. Similarly, this research revealed that the antifungal action of both lactic acid and acetic acid was enhanced by a decrease in the pH of the culture media (Vélez, C., León, A., 2014).

From the previous, it is suggested that an increase in the concentration of lactic acid will have a negative effect on the development of fungal colonies, thus inhibiting the presence of these microorganisms in the desired bacteriological culture medium. This concentration will only affect fungi up to a point, known as the minimum inhibitory concentration, which if exceeded would cause a concentration gradient of solutes between the internal and external medium of bacteria, which could cause an excessive outflow of water from these microorganisms, through osmotic movements, which would inhibit their growth.

### Purpose

During the last stage of education, normally the Baccalaureate, science subjects tend to emphasise the development of students’ practical or laboratory skills, where the aim is to put into practice the theoretical knowledge learned by the students. In Biology, one of the practical tasks that enriches students is the investigation of microorganisms, more specifically bacteria. Therefore, it is common that the development of culture media is a recurrent activity in the curricula, but, as a negative part of these explorations, these substrates are often contaminated, very commonly, by fungal colonies that not only prevent the correct control of variables but also inhibit the growth of bacteria; thus, preventing students from achieving the learning objectives of these practical lessons.

Due to this and the fact that commercial inhibitory media are often expensive, and their handling requires high complexity, the development of a fungal inhibitory medium using inexpensive agents was proposed. This will facilitate experimentation in microbiology at schools that do not have large laboratories and a large list of materials, thus contributing to the development of scientific knowledge among students globally and regardless of the resources they have, making science equitable and achievable.

## MATERIALS AND DEFINITION OF THE INVESTIGATION

### Variables

#### Dependent Variable

The dependent variable will be the number of bacterial and mycological colonies in the culture media, which will be measured using the quadrat technique. Subsequently, a statistical significance study will be carried to determine if there is a relationship between the lactic acid concentrations and the development of bacteria and fungi.

#### Controlled Variables

#### Independent Variable

The independent variable of this investigation is the lactic acid concentration (in %) in each culture media. This variable was modified in 5 different experimental groups, as it can be seen in table 1.

**Table 1.** Control and experimental groups of the research with their respective lactic acid concentrations (in %) in the culture media.

| *Group of the Investigation* | *Lactic Acid Concentration (in %)* |
| --- | --- |
| Control Group | 0 |
| Experimental Group 1 | 10 |
| Experimental Group 2 | 20 |
| Experimental Group 3 | 30 |
| Experimental Group 4 | 40 |
| Experimental Group 4 | 45 |

Due to the nature of this research, no controlled variables will be considered, as the purpose of this investigation is to perform a fungal inhibitory medium under any conditions, including the lack of standardisation in the methodologies.

### Materials

The required materials, dimensions, and uncertainties to carry out this investigation are reported in Table 2.

**Table 2.** Materials and equipment required together with their capacity and uncertainties (if applicable).

| *Materials* | *Measurement Equipment* | *Accessory Equipment* |
| --- | --- | --- |
| 1.05 L of distilled water.<br>30 grams of agar.<br>2 beef bouillon concentrate tablets Knorr Suiza.<br>100 ml of lactic acid solution with a concentration of 88%.<br>Solution containing 1 molar concentration of hydrochloric acid.<br>Solution containing 1 molar concentration of sodium hydroxide. | 2 test tubes with capacity of 50 ml and uncertainty of $\pm 0.5$ ml.<br>1 burette with capacity of 10 ml and uncertainty of $\pm 0.1$ ml.<br>1 beaker with capacity of 600 ml and uncertainty of $\pm 50.0$ ml.<br>9 beakers with capacity of 250 ml and uncertainty of $\pm 12.5$ ml.<br>1 dropper.<br>1 weighing scale with uncertainty of $\pm 0.0005$ g.<br>1 pH probe with uncertainty of $\pm 0.05$ units. | 1 hot plate.<br>1 magnetic stirrer.<br>1 crystallization dish with a capacity of 100 X 50.<br>6 glass stirrers.<br>3 metal spatulas.<br>1 watch glass.<br>24 petri dishes.<br>2 glass funnels with capacity of 100 mm.<br>24 swabs.<br>24 numbered labels.<br>2 latex gloves. |

### Methodology

#### Exploration A

Change in the concentration of lactic acid in the culture media solutions.

The aim of this exploration is to produce the solutions that will form the culture media. Each of the solutions will have a different concentration of lactic acid, in order to evaluate the antifungal role of this acid.

1. Using a volumetric measuring cylinder of 50 ml capacity, measure 300 ml of distilled water and place it in a 600 ml beaker.
2. Place the beaker with the distilled water on a hotplate and set a temperature of 400 ± 1 °C.
3. Bring the water to the boil and place the 1 tablet, 15 g of beef concentrate in the boiling water.
4. Place a magnetic stirrer in the beaker and switch on the stirrer magnet.
5. Once dissolved, add 6 grams of agar.
6. Bring the water to a temperature of 400 degrees, at a speed of 4 for 8 minutes.
7. Remove from the hotplate and leave to stand for 7 minutes.
8. Measure and adjust the pH to 7.5 units, using NaOH and HCl.
  a. If the pH needs to be reduced, add drops of HCl to the solution.
  b. If the pH is to be increased, add drops of NaOH to the solution.
9. In 6 beakers of 250 ml capacity dispense the lactic acid, using graduated cylinders and volumetric pipettes, according to the group as shown in Table 3.

**Table 3.**
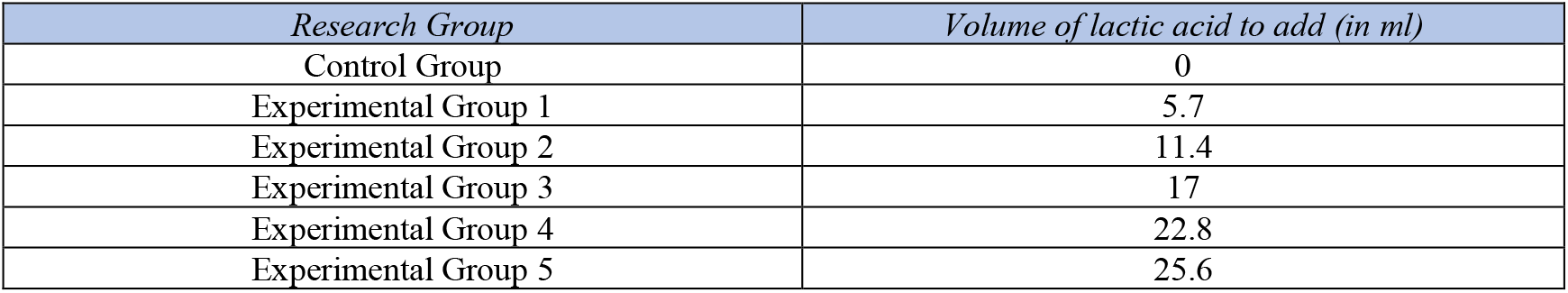
Volume of lactic acid (in ml) to be added to each solution by experimental and control group.
10. Add to each beaker the volume of agar and beef concentrated solution depending in the experimental or control group, as shown in table 4.

**Table 4.** Volume of agar and beef concentrated solution (in ml) to be added to each solution by experimental and control group.
11. Using a separate glass stirrer for each group, stir the solutions.
12. Make 4 cultures per experimental group, i.e. a total of 24 petri dishes including repetitions and the control group shall be used.
13. Using labels, label according to experimental group as follows:

| *Research Group* | *Volume of agar and beef concentrated solution to add (in ml)* |
| --- | --- |
| Control Group | 50 |
| Experimental Group 1 | 44.3 |
| Experimental Group 2 | 38.6 |
| Experimental Group 3 | 33 |
| Experimental Group 4 | 27.3 |
| Experimental Group 5 | 24.4 |

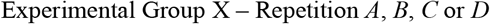
14. Once the petri dishes with the cultures have been prepared, let the cultures stand for 8 days; to achieve a gel consistency in the media due to the effect of the agar.

#### Exploration B

Seeding of the culture media.

The purpose of this exploration is to carry out microbiological seeding on the culture media of the two scans, to evaluate the antifungal potentials of the independent variables.

1. Using latex gloves, obtain a saliva sample by inserting a swab into the oral cavity.
2. Once the sample has been obtained, “draw” on the culture media, produced in exploration A, with the swab containing the sample.
3. Incubate the samples for four days.
4. After seeding is complete, discard the swabs in a waste bag. Perform steps 1 to 4 for each replicate of each experimental group.

#### Exploration C

Observation and data recording.

Perform an observation, using the quadrat technique, of fungal and bacterial colonies developed on the media of examination A.

### Justification

In the exploration A, lactic acid was chosen for its effect on the electrochemical gradient of the cell membrane. On the other hand, the volumes of this acid were arranged according to the desired concentration of each experimental group, so that it would affect the fungi but not the bacteria, thus providing a bacteriological specific medium; these volumes were calculated as described in appendix 1.

## RESULTS

### Qualitative Data

After analysing the culture media, no qualitative data was found that could affect, either positively or negatively, the purposes of this research.

### Quantitative Data

For bacterial and fungal colony counts, the data shown in tables 5-6 were obtained.

**Table 5.** Number of fungal populations by experimental and control groups and their repetitions with lactic acid concentrations of 0, 10, 20, 30, 40 y 45%.

| **Number of Fungal Populations** |  |  |  |  |
| --- | --- | --- | --- | --- |
| *Experimental Group* | *Repetition A* | *Repetition B* | *Repetition C* | *Repetition D* |
| Control Group<br>(Lactic acid concentration of 0%) | 8 | 7 | 10 | 12 |
| E. G. 1<br>(Lactic acid concentration of 10%) | 0 | 0 | 0 | 0 |
| E. G. 2<br>(Lactic acid concentration of 20%) | 0 | 0 | 0 | 0 |
| E. G. 3<br>(Lactic acid concentration of 30%) | 0 | 0 | 0 | 0 |
| E. G. 4<br>(Lactic acid concentration of 40%) | 0 | 0 | 0 | 0 |
| E. G. 5<br>(Lactic acid concentration of 45%) | 0 | 0 | 0 | 0 |

**Table 6.** Number of bacterial populations by experimental and control groups and their repetitions with lactic acid concentrations of 0, 10, 20, 30, 40 y 45%.

| Number of Bacterial Populations |  |  |  |  |
| --- | --- | --- | --- | --- |
| Experimental Group | Repetition A | Repetition B | Repetition C | Repetition D |
| Control Group<br>(Lactic acid concentration of 0%) | 0 | 0 | 0 | 0 |
| E. G. 1<br>(Lactic acid concentration of 10%) | 12 | 9 | 13 | 10 |
| E. G. 2<br>(Lactic acid concentration of 20%) | 14 | 15 | 17 | 15 |
| E. G. 3<br>(Lactic acid concentration of 30%) | 13 | 10 | 11 | 9 |
| E. G. 4<br>(Lactic acid concentration of 40%) | 7 | 10 | 9 | 9 |
| E. G. 5<br>(Lactic acid concentration of 45%) | 8 | 6 | 5 | 5 |

### Processed Data

The averages of bacterial and fungal colonies were calculated, in tables 7-8, to obtain a mean value of the number of colonies that will facilitate data management. Also, the standard deviations were calculated, as shown in appendix 2, for each data set, i.e. for each experimental group and its repetitions, which will allow to analyse the precision and accuracy of the investigation.

**Table 7.** The final mean number of fungal populations per experimental group and the standard deviation of this mean are shown.

| Number of Fungal Populations |  |  |  |  |  |  |
| --- | --- | --- | --- | --- | --- | --- |
| Experimental Group | Repetition A | Repetition B | Repetition C | Repetition D | Average | Standard Deviation |
| Control Group<br>(Lactic acid concentration of 0%) | 8 | 7 | 10 | 12 | 9.25 | 3.68 |
| E. G. 1<br>(Lactic acid concentration of 10%) | 0 | 0 | 0 | 0 | 0 | 0 |
| E. G. 2<br>(Lactic acid concentration of 20%) | 0 | 0 | 0 | 0 | 0 | 0 |
| E. G. 3<br>(Lactic acid concentration of 30%) | 0 | 0 | 0 | 0 | 0 | 0 |
| E. G. 4<br>(Lactic acid concentration of 40%) | 0 | 0 | 0 | 0 | 0 | 0 |
| E. G. 5<br>(Lactic acid concentration of 45%) | 0 | 0 | 0 | 0 | 0 | 0 |

**Table 8.** The final mean number of bacterial populations per experimental group and the standard deviation of this mean are shown.

| Number of Bacterial Populations |  |  |  |  |  |  |
| --- | --- | --- | --- | --- | --- | --- |
| Grupo experimental | Repetition A | Repetition B | Repetition C | Repetition D | Average | Standard Deviation |
| Control Group (Lactic acid concentration of 0%) | 0 | 0 | 0 | 0 | 0 | 0 |
| E. G. 1 (Lactic acid concentration of 10%) | 12 | 9 | 13 | 10 | 11 | 2.5 |
| E. G. 2 (Lactic acid concentration of 20%) | 14 | 15 | 17 | 15 | 15.25 | 1.18 |
| E. G. 3 (Lactic acid concentration of 30%) | 13 | 10 | 11 | 9 | 10.75 | 2.19 |
| E. G. 4 (Lactic acid concentration of 40%) | 7 | 10 | 9 | 9 | 8.75 | 1.19 |
| E. G. 5 (Lactic acid concentration of 45%) | 8 | 6 | 5 | 5 | 6 | 1.5 |

### Statistical Analysis

A t-test was performed, which allows to know statistically whether the dependent variable is significant with the independent variable, according to the degrees of freedom and statistical significance shown in appendix 3.

To perform this test, it is necessary to establish the null and alternative hypotheses of the research, as shown in table 9.

**Table 9.** The null and alternative hypothesis of the research are shown and explained.

| Null Hypothesis | Alternative Hypothesis |
| --- | --- |
| Lactic acid will not have any significance for fungal growth in the culture media. | Lactic acid will have a statistical significance on fungal growth in culture media; either positively or negatively. |

**Table 10.** The degrees of freedom and its meaning is shown in relation with the T-test.

| Degrees of Freedom | Meaning |
| --- | --- |
| 2.01 | If the T-test coefficient exceeds the degrees of freedom, the alternative hypothesis is approved; if a lower value is obtained, the null hypothesis is approved. |

**Table 11.** The values obtained from the T-test, the critical value, and the comparison between these values for the approval or rejection of the hypotheses are shown.

| Correlation | T-test coefficient | Critical Value | Approved Hypothesis |
| --- | --- | --- | --- |
| Lactic acid and fungal populations | 3.1 | 2.01 | Alternative hypothesis |
| Lactic acid and bacterial populations | 2.1 | 2.01 | Alternative hypothesis |

Subsequently, the α value was established, which represents the level of reliability necessary to draw conclusions correctly, which is 0.05, meaning that the conclusions drawn will have 95% validity or reliability; this makes it possible to establish what is shown in table 15 according to the nature of each experiment.

The t-test was performed by means of the data processor PLANTECALC, which will obtain the coefficient that will later be compared with the established degrees of freedom, as shown in table (PlanetCALC., n.d.).

### Presentation

Figure 1 shows the processed data in the shape of a polynomic line.

**Figure 1.**
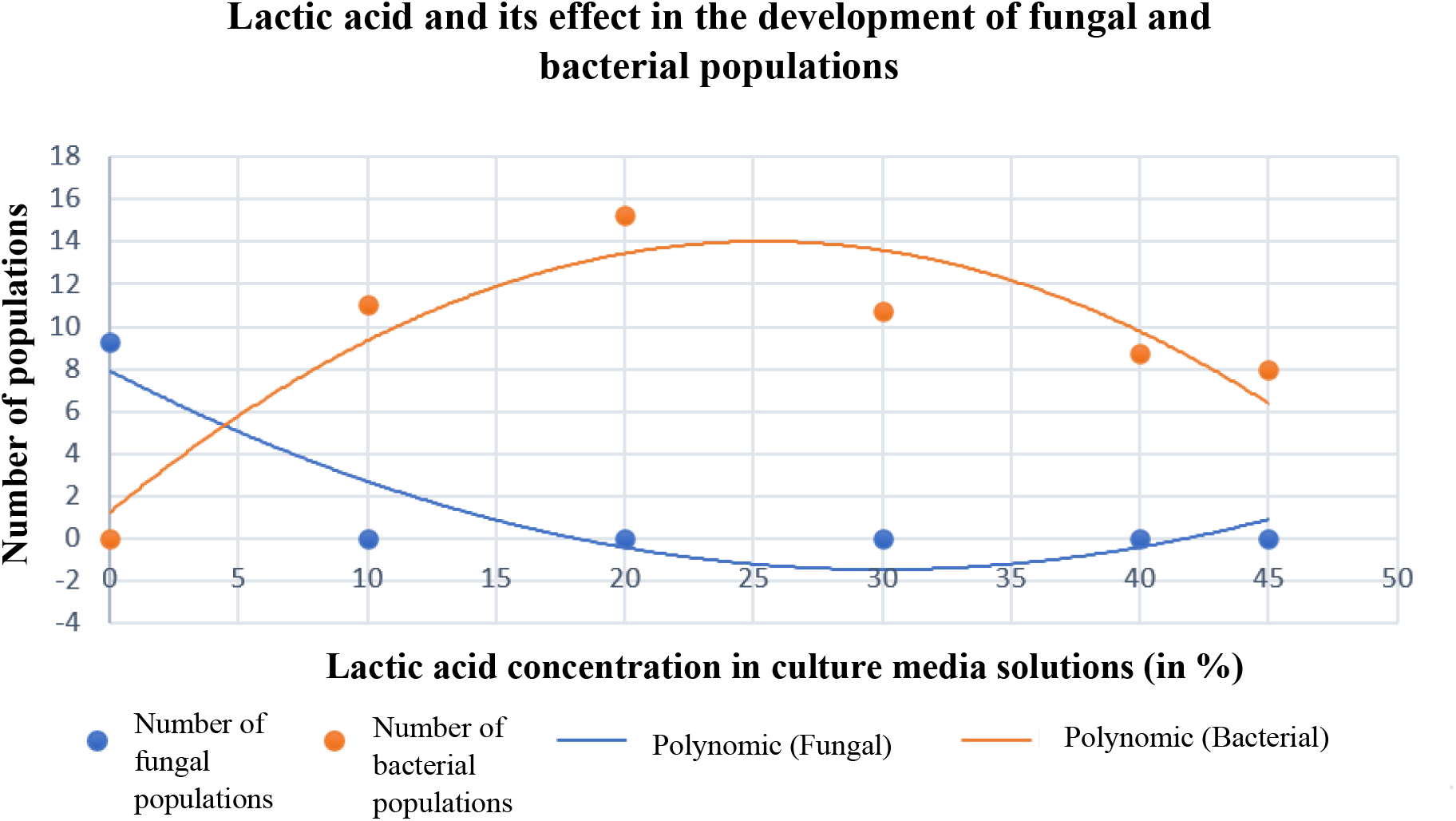
The correlation between lactic acid concentration (in %) and the number of bacterial and fungal populations is shown.

## DISCUSSION

Finally, an analysis and interpretation of the data obtained from experiment A will be carried out, from which the role of lactic acid as an antifungal agent will be defined.

Regarding the statistical analysis carried out, it is possible to conclude the following:

I. There is a correlation between the lactic acid concentrations of the media and the number of fungal colonies; the alternative hypothesis is approved.
II. Similarly, the alternative hypothesis between lactic acid concentration and bacterial growth was approved; the alternative hypothesis was approved.

In fungal growth, an inversely proportional relationship with lactic acid concentration can be established. This is because the statistical analysis concluded a relationship of significance between these two variables; subsequently, and thanks to the modelling of the data, it is observed how an increase in the concentration of lactic acid produces a decrease in the number of colonies. In the same way, this can be supported by external literature, which affirms the role of lactic acid as a neutralising agent of the electrochemical gradient of fungal membranes, which prevents cell transport and, therefore, inhibits the development of these microorganisms (Vélez, C., León, A., 2014).

On the other hand, the study of the relationship between lactic acid and the number of bacterial colonies shows a significance that, as shown in graph 1, has a double behaviour. Firstly, in the concentrations from 0 to 20%, a positive correlation is observed, meaning that an increase in the concentration of lactic acid increases bacterial development. in contrast, from this last concentration onwards, an inverse correlation between the variables is observed, meaning that an increase in the concentration produces a decrease in bacterial growth.

Interpreting these trends, it can be argued that the positive relationship between bacteria and lactic acid is due to the fact that these microorganisms take advantage of the compounds present in the acid for their metabolism. This utilization would only be seen up to a concentration of 20%, where a negative correlation would be present. This is due to the concentration gradient that is established between the cellular and external environment, which would lead to the dehydration of the bacteria by the outflow of water to the external environment in order to achieve an isotonic solution in which the solute/solvent balance is the same in the internal (cellular) and external environments.

### Evaluation of the Methodology

For the purpose of this research, the use of controlled variables was not necessary, as the aim was to develop a fungal inhibitory culture medium under all conditions, even without following a standardised methodology.

In an analysis of the methodology, it was chosen and practiced correctly, as all the steps were studied during the preliminary investigations and explorations, which allowed to know the impact that the methodology was going to have on the results of the research.

Regarding the reliability of the methodology, results, and conclusions, it can be said that this would have been increased if more replications per experimental group had been used. This is because the possibility of random errors affecting the results would be reduced.

Finally, ethical, health, and environmental considerations were taken into account throughout the experimentation. This is because all scientific research must consider aspects that may not have a direct impact on the research but could have an impact on the context in which it is carried out, such as water waste or poor handling of microorganisms.

### Result’s Assessment

As can be seen in the results, no errors, neither random nor standardised, were found in the research, which, together with the analysis of the methodology, gives the conclusions drawn a high level of reliability.

Finally, thanks to the statistical analysis carried out, the conclusions drawn have a high level of reliability, as a standard of 95% reliability was established. This not only demonstrates reliability in a statistical way but is also accompanied by the interpretation of the data; thus, explaining the conclusions drawn in a mathematical and biological way.

### Criteria for Literature Inclusion

The literature inclusion criteria were delimited in such a way that the bibliographic sources used could be contrasted with other sources, except for the case of the role of lactic acid as an antifungal due to the absence of previous research. Also, the bibliography was compiled from reliable sources, such as universities or popular articles.

## CONCLUSIONS

It is concluded that lactic acid could inhibit fungal growth. This effect, if bacterial growth is desired, will be achieved at a concentration of this acid of 20%, this being the minimum inhibitory concentration, since otherwise bacterial growth would be impaired.

Thanks to this research, it will be possible to develop, in school contexts or in those where resources are not available, inhibitory means that will allow a better study of bacteria and their behaviour without contamination by fungi.

Finally, the application of this novel technique will not only be useful in the aforementioned contexts, but it is also possible to foresee an application in laboratories, where high technology is not available, for the development of research, since, in the same way as sophisticated and expensive inhibitory culture media, this culture medium is capable of inhibiting fungal growth and, to a lesser extent, promoting bacterial development; proving the use of lactic acid as an antifungal agent.

## SUPPLEMENTARY INFORMATION

Supplementary information on this work can be found in the section called “appendices”, where raw data, calculations, and statistical tables can be found for a complete understanding of this paper.

## ACKNOWLEDGEMENTS

From this project, I would like to thank and acknowledge the special and fundamental importance in the development of this research to the Biology teacher, Diana Guzmán, and the Laboratory Assistant, Patricia Galindo, from PrepaTEC campus Santa Fe; as well as to all the directors and coordinators who made possible the use of the laboratory and other resources of this campus.

## APPENDICES Appendix 1

This appendix shows the calculations made to obtain the amount of lactic acid required, per experimental group, according to the desired concentration.

|  |  |  |
| --- | --- | --- |
| **Concentration Formula**<br>$V_1C_1 = V_2C_2$ | Solve for $V_1$ | $V_1 = V_2 \times C_2 / C_1$ |

| Calculation |  |  |  |  |  |  |
| --- | --- | --- | --- | --- | --- | --- |
| Groups | $V_1$ | $C_1$ | $V_2$ | $C_2$ | Multiplication | Division |
| GC: | ? | 88 | 50 | 0 | 0 | 0 |
| G.E.1: | ? | 88 | 50 | 10 | 500 | 5,68181818 |
| G.E.2: | ? | 88 | 50 | 20 | 1000 | 11,3636364 |
| G.E.3: | ? | 88 | 50 | 30 | 1500 | 17,0454545 |
| G.E.4: | ? | 88 | 50 | 40 | 2000 | 22,7272727 |
| G.E.5: | ? | 88 | 50 | 45 | 2250 | 25,5681818 |

| Groups | $V_2$ | $V_1$ | Volume of the media solution |
| --- | --- | --- | --- |
| GC: | 50 | 0 | 50 |
| G.E.1: | 50 | 5,7 | 44,3 |
| G.E.2: | 50 | 11,36 | 38,64 |
| G.E.3: | 50 | 17,04 | 32,96 |
| G.E.4: | 50 | 22,72 | 27,28 |
| G.E.5: | 50 | 25,57 | 24,43 |

## Appendix 2

The standard deviation of each experimental and control group was calculated with the following formula:

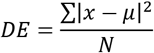

Where:

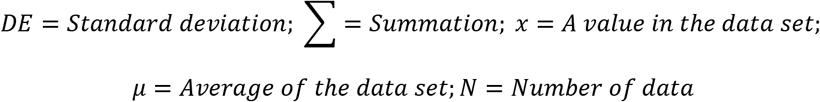

With this formula, the following tables were constructed with the data processer *Excel* in order to calculate the standard deviation:

### Appendix 2.1

Calculation of the standard deviation of the fungal populations that were placed in culture media with different lactic acid concentrations.

| Control Group |  |  |  |  |  |  |
| --- | --- | --- | --- | --- | --- | --- |
| Repetition | Value | Average | Subtraction | Rest squared | Addition | Division |
| Repetition A | 8 | 9,25 | -1,25 | 1,5625 | 14,75 | 3,6875 |
| Repetition B | 7 | 9,25 | -2,25 | 5,0625 |  |  |
| Repetition C | 10 | 9,25 | 0,75 | 0,5625 |  |  |
| Repetition D | 12 | 9,25 | 2,75 | 7,5625 |  |  |

| Experimental Group 1 |  |  |  |  |  |  |
| --- | --- | --- | --- | --- | --- | --- |
| Repetition | Value | Average | Subtraction | Rest squared | Addition | Division |
| Repetition A | 0 | 0 | 0 | 0 | 0 | 0 |
| Repetition B | 0 | 0 | 0 | 0 |  |  |
| Repetition C | 0 | 0 | 0 | 0 |  |  |
| Repetition D | 0 | 0 | 0 | 0 |  |  |

| Experimental Group 2 |  |  |  |  |  |  |
| --- | --- | --- | --- | --- | --- | --- |
| Repetition | Value | Average | Subtraction | Rest squared | Addition | Division |
| Repetition A | 0 | 0 | 0 | 0 | 0 | 0 |
| Repetition B | 0 | 0 | 0 | 0 |  |  |
| Repetition C | 0 | 0 | 0 | 0 |  |  |
| Repetition D | 0 | 0 | 0 | 0 |  |  |

| Experimental Group 3 |  |  |  |  |  |  |
| --- | --- | --- | --- | --- | --- | --- |
| Repetition | Value | Average | Subtraction | Rest squared | Addition | Division |
| Repetition A | 0 | 0 | 0 | 0 | 0 | 0 |
| Repetition B | 0 | 0 | 0 | 0 |  |  |
| Repetition C | 0 | 0 | 0 | 0 |  |  |
| Repetition D | 0 | 0 | 0 | 0 |  |  |

| Experimental Group 4 |  |  |  |  |  |  |
| --- | --- | --- | --- | --- | --- | --- |
| Repetition | Value | Average | Subtraction | Rest squared | Addition | Division |
| Repetition A | 0 | 0 | 0 | 0 | 0 | 0 |
| Repetition B | 0 | 0 | 0 | 0 |  |  |
| Repetition C | 0 | 0 | 0 | 0 |  |  |
| Repetition D | 0 | 0 | 0 | 0 |  |  |

| Experimental Group 5 |  |  |  |  |  |  |
| --- | --- | --- | --- | --- | --- | --- |
| Repetition | Value | Average | Subtraction | Rest squared | Addition | Division |
| Repetition A | 0 | 0 | 0 | 0 | 0 | 0 |
| Repetition B | 0 | 0 | 0 | 0 |  |  |
| Repetition C | 0 | 0 | 0 | 0 |  |  |
| Repetition D | 0 | 0 | 0 | 0 |  |  |

### Appendix 2.2

Calculation of the standard deviation of the bacterial populations that were placed in culture media with different lactic acid concentrations.

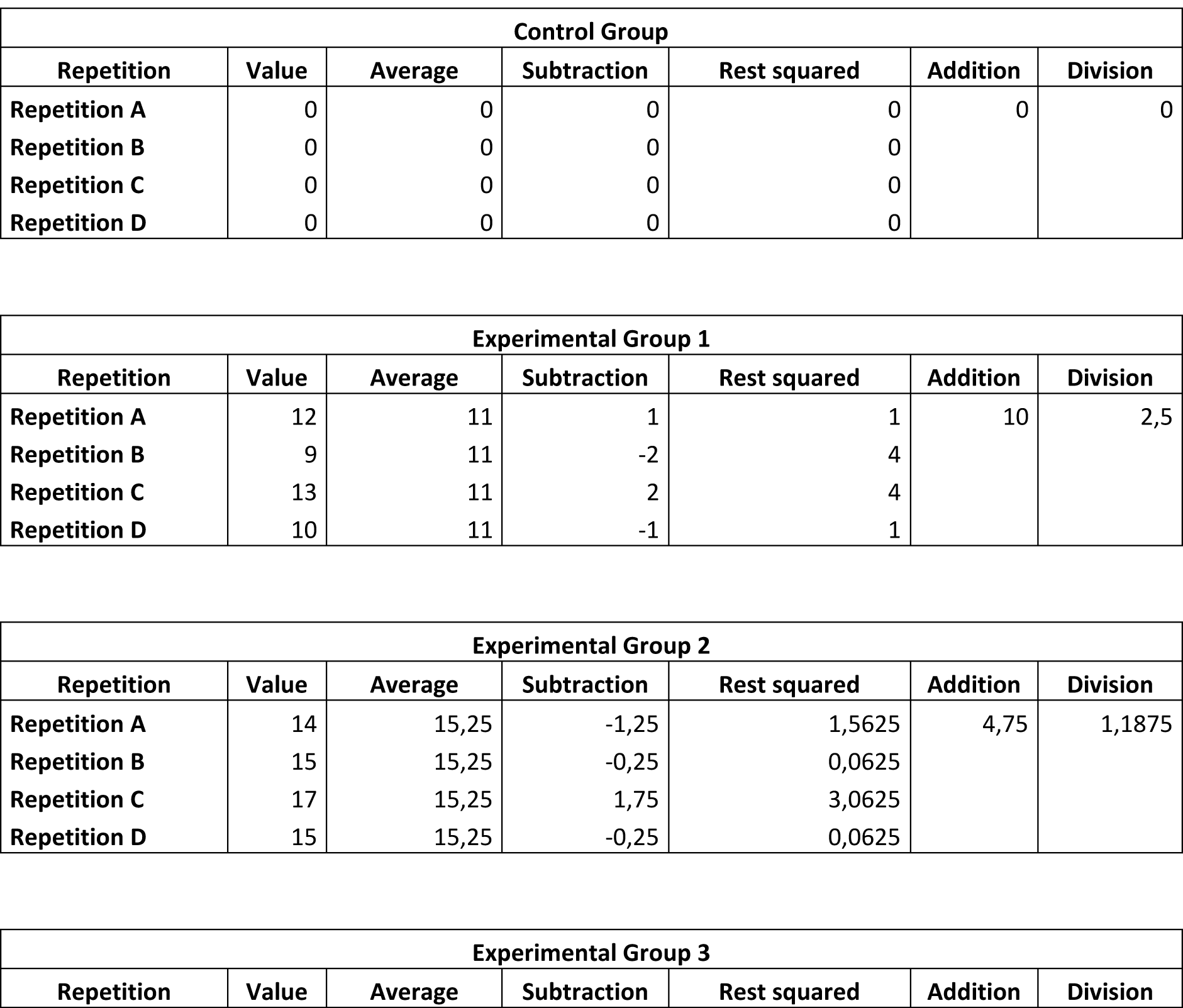
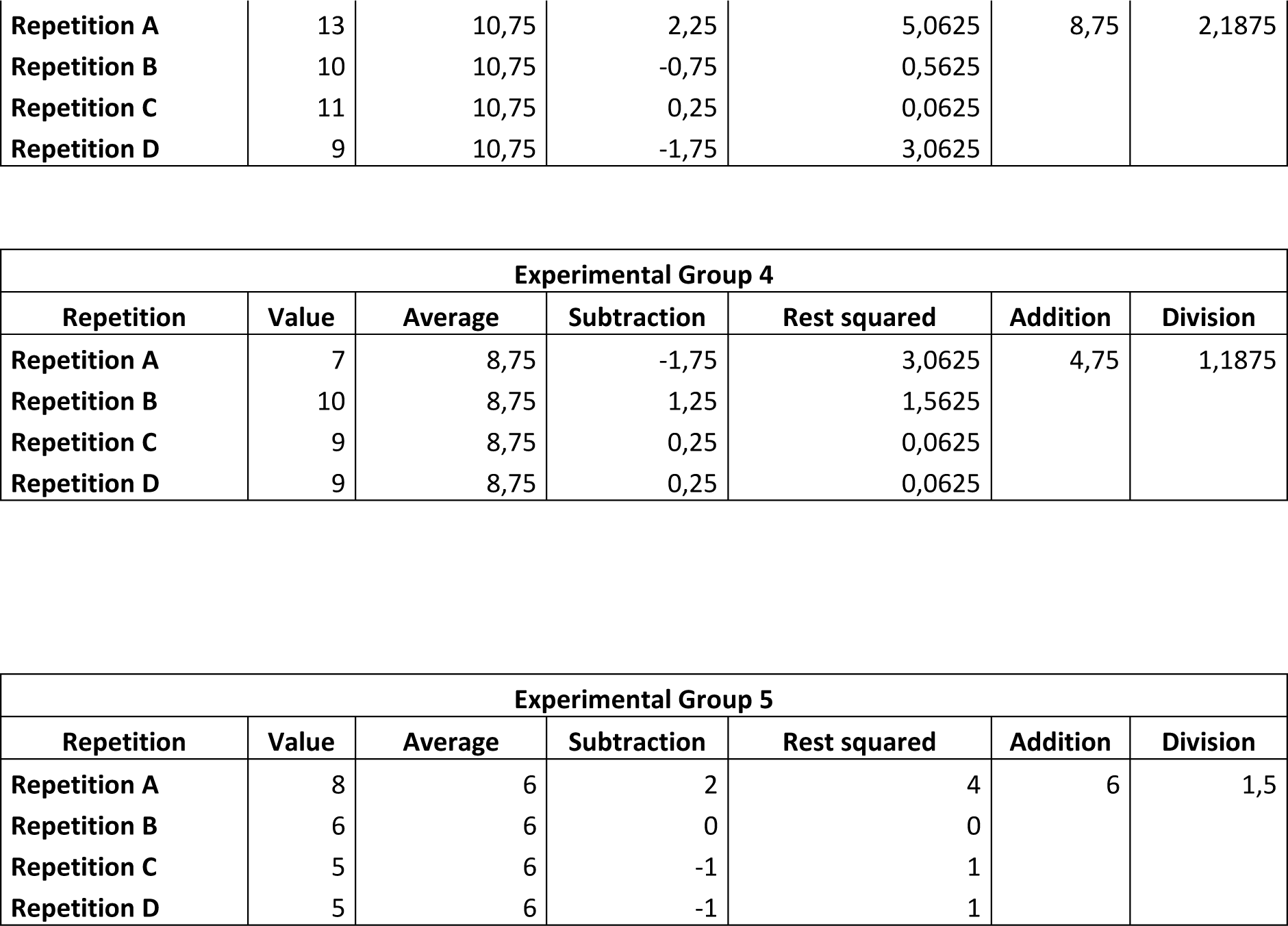

| Control Group |  |  |  |  |  |  |
| --- | --- | --- | --- | --- | --- | --- |
| Repetition | Value | Average | Subtraction | Rest squared | Addition | Division |
| Repetition A | 0 | 0 | 0 | 0 | 0 | 0 |
| Repetition B | 0 | 0 | 0 | 0 |  |  |
| Repetition C | 0 | 0 | 0 | 0 |  |  |
| Repetition D | 0 | 0 | 0 | 0 |  |  |

| Experimental Group 1 |  |  |  |  |  |  |
| --- | --- | --- | --- | --- | --- | --- |
| Repetition | Value | Average | Subtraction | Rest squared | Addition | Division |
| Repetition A | 12 | 11 | 1 | 1 | 10 | 2,5 |
| Repetition B | 9 | 11 | -2 | 4 |  |  |
| Repetition C | 13 | 11 | 2 | 4 |  |  |
| Repetition D | 10 | 11 | -1 | 1 |  |  |

| Experimental Group 2 |  |  |  |  |  |  |
| --- | --- | --- | --- | --- | --- | --- |
| Repetition | Value | Average | Subtraction | Rest squared | Addition | Division |
| Repetition A | 14 | 15,25 | -1,25 | 1,5625 | 4,75 | 1,1875 |
| Repetition B | 15 | 15,25 | -0,25 | 0,0625 |  |  |
| Repetition C | 17 | 15,25 | 1,75 | 3,0625 |  |  |
| Repetition D | 15 | 15,25 | -0,25 | 0,0625 |  |  |

| Experimental Group 3 |  |  |  |  |  |  |
| --- | --- | --- | --- | --- | --- | --- |
| Repetition | Value | Average | Subtraction | Rest squared | Addition | Division |
| Hinojosa. – Lactic acid as an antifungal factor |  |  |  |  |  |  |
| **Repetition A** | 13 | 10,75 | 2,25 | 5,0625 | 8,75 | 2,1875 |
| **Repetition B** | 10 | 10,75 | -0,75 | 0,5625 |  |  |
| **Repetition C** | 11 | 10,75 | 0,25 | 0,0625 |  |  |
| **Repetition D** | 9 | 10,75 | -1,75 | 3,0625 |  |  |

| **Experimental Group 4** |  |  |  |  |  |  |
| --- | --- | --- | --- | --- | --- | --- |
| **Repetition** | **Value** | **Average** | **Subtraction** | **Rest squared** | **Addition** | **Division** |
| **Repetition A** | 7 | 8,75 | -1,75 | 3,0625 | 4,75 | 1,1875 |
| **Repetition B** | 10 | 8,75 | 1,25 | 1,5625 |  |  |
| **Repetition C** | 9 | 8,75 | 0,25 | 0,0625 |  |  |
| **Repetition D** | 9 | 8,75 | 0,25 | 0,0625 |  |  |

| **Experimental Group 5** |  |  |  |  |  |  |
| --- | --- | --- | --- | --- | --- | --- |
| **Repetition** | **Value** | **Average** | **Subtraction** | **Rest squared** | **Addition** | **Division** |
| **Repetition A** | 8 | 6 | 2 | 4 | 6 | 1,5 |
| **Repetition B** | 6 | 6 | 0 | 0 |  |  |
| **Repetition C** | 5 | 6 | -1 | 1 |  |  |
| **Repetition D** | 5 | 6 | -1 | 1 |  |  |

## Appendix 3

The following image demonstrates the relationship between the critical values and the degrees of freedom necessary for the correct interpretation of the t-test, performed as the statistical analysis of this research.

**Figure 2.**
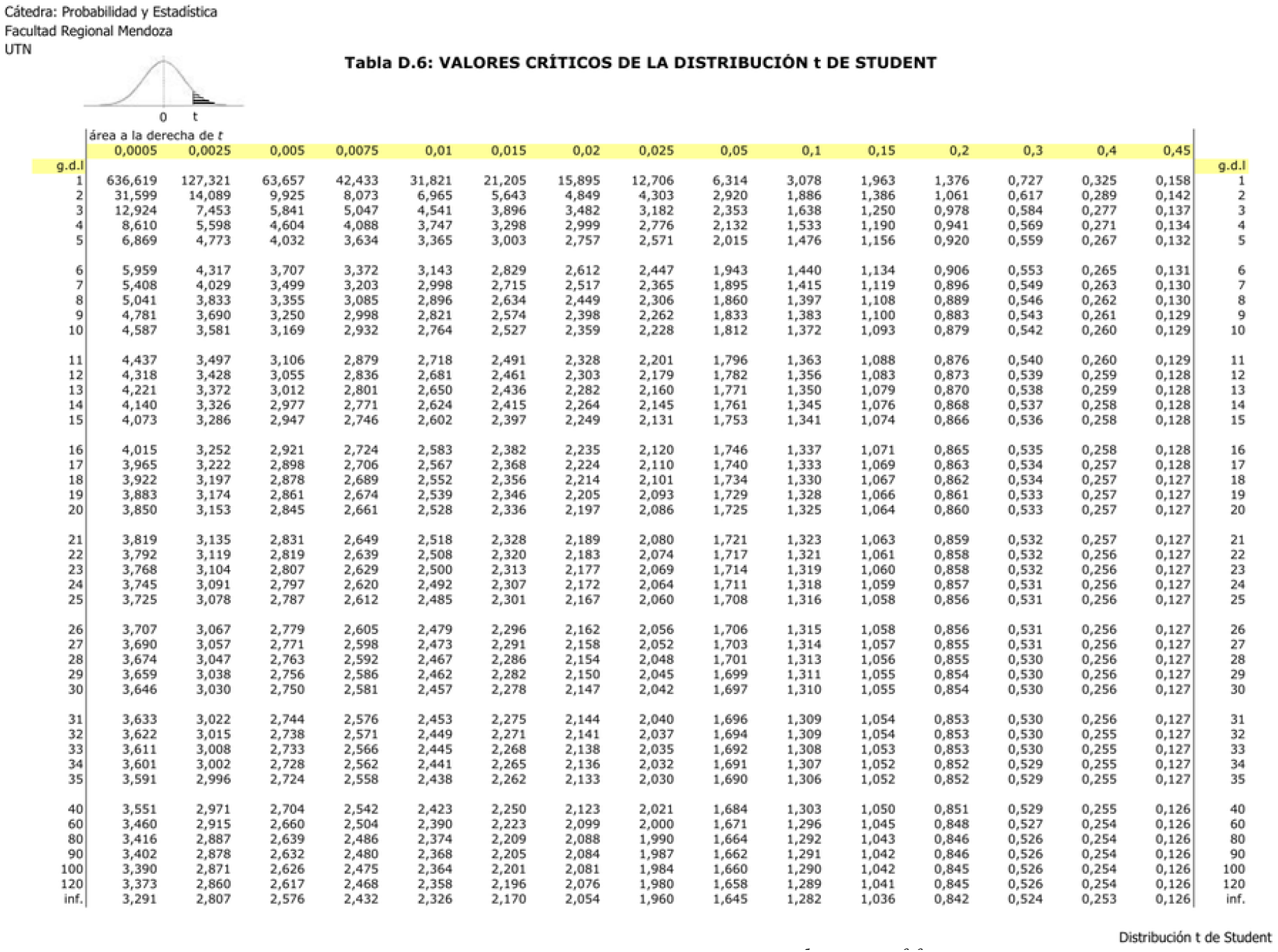
The table of critical values and degrees of freedom for their distribution in the t-test is shown. (Facultad Regional Mendoza., n.d.)

